# The Immune Signatures Data Resource: A compendium of systems vaccinology datasets

**DOI:** 10.1101/2021.11.05.465336

**Authors:** Joann Diray-Arce, Helen E.R. Miller, Evan Henrich, Bram Gerritsen, Matthew P. Mulè, Slim Fourati, Jeremy Gygi, Thomas Hagan, Lewis Tomalin, Dmitry Rychkov, Dmitri Kazmin, Daniel G. Chawla, Hailong Meng, Patrick Dunn, John Campbell, The Human Immunology Project Consortium (HIPC), Minnie Sarwal, John S. Tsang, Ofer Levy, Bali Pulendran, Rafick Sekaly, Aris Floratos, Raphael Gottardo, Steven H. Kleinstein, Mayte Suárez-Fariñas

## Abstract

Vaccines are among the most cost-effective public health interventions for preventing infection-induced morbidity and mortality, yet much remains to be learned regarding the mechanisms by which vaccines protect. Systems immunology combines traditional immunology with modern ‘omic profiling techniques and computational modeling to promote rapid and transformative advances in vaccinology and vaccine discovery. The NIH/NIAID Human Immunology Project Consortium (HIPC) has leveraged systems immunology approaches to identify molecular signatures associated with the immunogenicity of many vaccines, including those targeting seasonal influenza, yellow fever, and hepatitis B. These data are made available to the broader scientific community through the *ImmuneSpace* data portal and analysis engine leveraging the NIH/NIAID *ImmPort* repository^1,2^. However, a barrier to progress in this area is that comparative analyses have been limited by the distributed nature of some data, potential batch effects across studies, and the absence of multiple relevant studies from non-HIPC groups in *ImmPort.* To support comparative analyses across different vaccines, we have created the Immune Signatures Data Resource, a compendium of standardized systems vaccinology datasets. This data resource is available through *ImmuneSpace*, along with code to reproduce the processing and batch normalization starting from the underlying study data in *ImmPort* and the Gene Expression Omnibus (GEO). The current release comprises 1405 participants from 53 cohorts profiling the response to 24 different vaccines and includes transcriptional profiles and antibody response measurements. This novel systems vaccinology data release represents a valuable resource for comparative and meta-analyses that will accelerate our understanding of mechanisms underlying vaccine responses.

|  |  |
| --- | --- |
| Design Type(s) | ☐ Factorial design<br>☐ Longitudinal design<br>☐ Transcription profiling design |
| Factor Type(s) | ☐ Sample type (PBMC, Blood)<br>☐ Vaccine<br>☐ Pathogen<br>☐ Age groups |
| Measurement Type(s) | ☐ Transcription profiling assay<br>☐ Neutralizing antibody assay<br>☐ ELISA<br>☐ Influenza hemagglutination inhibition assay (HAI) |
| Technology Type(s) | ☐ Microarray<br>☐ RNA sequencing |
| Sample Characteristic(s) | ☐ Homo sapiens |

## Background and Summary

Vaccines, one of humanity’s greatest public health achievements, save millions of lives every year by preventing infectious diseases^3,4^. Despite their widespread use and efficacy, much remains to be learned regarding their molecular mechanisms of action. This is true both for vaccines against pandemic infections such as influenza^5^, and SARS-coronavirus-2^6^, as well as for infections for which there are currently no authorized or approved vaccines such as HIV^7–9^. Elucidating the commonalities and differences in the immune responses induced by different vaccines and their association with protective antibody responses will provide deeper insight and a framework for the evidence-based design of better vaccines or vaccination strategies. Recent technologies have provided tools to probe the immune response to vaccination and integrate hierarchical levels of the biological system^10^. Alluded to as systems vaccinology^11^, this new application of systems biology tools provides new insights into molecular mechanisms of vaccine-induced immunogenicity and protection^12–15^.

The National Institute of Allergy and Infectious Diseases (NIAID) established a multi-institutional consortium, Human Immunology Project Consortium (HIPC)^2,16^, to characterize the immune system in diverse populations in response to a stimulus, such as vaccination, using high-dimensional ‘omic platforms and modern computational tools^2^. Since the inception of the consortium in 2010, members of HIPC have published > 500 articles, including many that describe molecular signatures associated with vaccine-induced protection. These studies include molecular signatures that predict the immunogenicity of vaccination against yellow fever^17–20^, seasonal influenza in healthy young adults, elderly^21–25^, and children^26^, shingles^27,28^, dengue^29,30^, malaria^31,32^, and meta-analyses of common signatures across different vaccines^33,34^. These molecular signatures resulted from large-scale data analysis using high-throughput systems biology approaches coupled with detailed clinical phenotyping in well-characterized human cohorts.

Predicting immunogenicity from ‘omic signatures remains challenging, prompting methodological innovation to advance the field towards delivering on the promises of precision vaccination^35–37^. The factors that contribute to robust vaccination responses are highly complex and span multiple biological scales. The vast collection of high-dimensional profiling datasets poses significant challenges for comparative analysis of these studies, including biological variability as well as data challenges such as volume, technical noise, and diverse sample processing pipelines. Data integration of cellular and molecular signatures to predict vaccine responses requires harmonization and normalization of data from multiple sources^38^. The generation of big data poses simultaneous challenges and opportunities with the potential of contributing to precision medicine. The biological interpretation of the resulting molecular features correlated with robust responses is another key factor. Understanding how effective vaccines stimulate protective immune responses, and how these mechanisms may differ between vaccine types and targeted pathogens remains a substantial challenge for the field. Moreover, the systems vaccinology field has been limited by a lack of a formal framework to standardize immune signatures gathered from diverse studies, creating a bottleneck for comparative analysis. To address these challenges, and in support of advances in systems vaccinology by the HIPC project and the broader scientific community, we present the creation of the Immune Signatures Data Resource, a compendium of systems vaccinology studies that enables standardized comparative analysis to identify molecular signatures that correlate with the outcomes of vaccinations.

The current release of the Immune Signatures Data Resource consists of 4795 transcriptomic samples from 1405 participants curated from 30 *ImmPort* studies (16 from HIPC-related studies, 14 non-HIPC studies) (Figure 2). The transcriptomic profiling dataset is derived from 53 cohorts of 820 young adults (18-49 years old) and 585 (≥50 years old) older adult samples. The data resource covers 24 vaccines targeting 11 pathogens and 6 vaccine types (Figure 1B, 4A), thus creating a critical mass of data that will serve as a valuable resource for the broader scientific community. Additionally, data assembly and integration of these data set enables derivation of comparable signatures for each study for comparative analysis of the underlying data.

**Figure 1:**
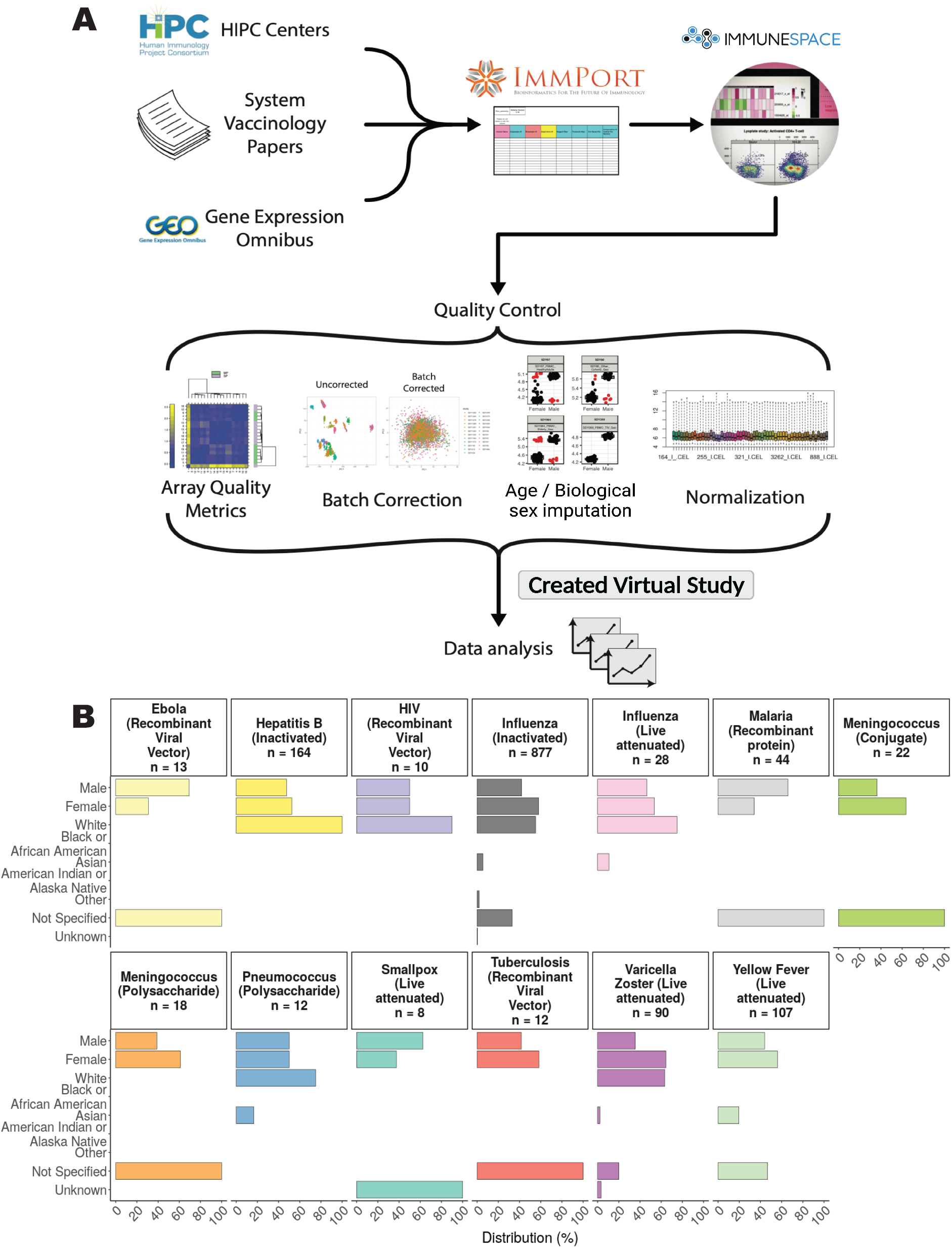
HIPC Immune Signatures Data Resource pipeline and study demographics. A. Systems vaccinology datasets from existing HIPC studies, as well as published systems vaccinology papers and databases, were submitted to the *ImmPort* database. *ImmuneSpace* captures these datasets to create a combined compendium dataset. Quality control assessments of these data include array quality checks for microarray studies, batch correction, imputations for missing age and sex/y-chromosome presence information, and normalization per study. The combined virtual study included transcriptional profiles and antibody response measurements from 1405 participants across 53 cohorts, profiling the response to 24 different vaccines. B. Demographic data included biological sex, race, vaccine, and number of participants.

**Figure 2:**
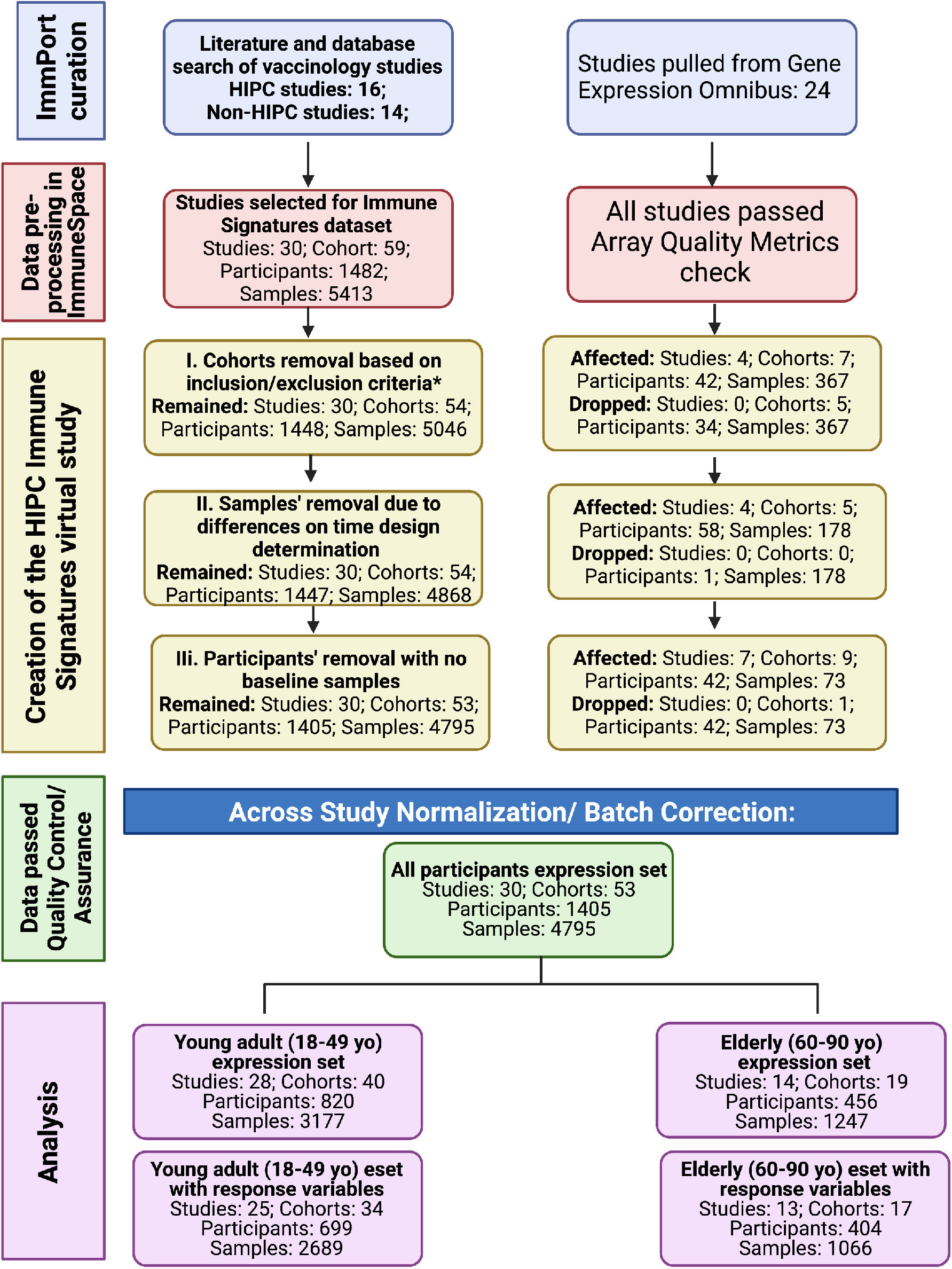
Flow chart diagram of the Immune Signatures Data Resource.

## Methods

### Database background information and structure

#### Compatibility with ImmPort and ImmuneSpace, the central databases of the Human Immunology Project Consortium

Given the exponential growth of the number of datasets of multiple modalities, an urgent need emerged for data sharing across the broader scientific community. The HIPC implements the NIH Data Sharing policy to promote the principles of Findability, Accessibility, Interoperability, and Reusability (FAIR) via *ImmPort*, created under the National Institute of Allergy and Infectious Diseases Division of Allergy, Immunology, and Transplantation (NIAID-DAIT). *ImmPort* (ImmPort.org) is an open repository of participant-level large-scale human immunology data designed to aid scientists with data standards and guidelines for efficient secondary analyses^1,39^. *ImmPort* facilitates data sharing of immunology studies creating a centralized knowledge base and resources, and serves as a central data repository for HIPC. *ImmuneSpace*^2,34^ extends *ImmPort*, providing access to additional data (e.g., standardized gene expression matrices) and web-based R tools for data accession, analysis, and reporting. Studies in the Immune Signatures Data Resource are archived through the Shared Data Portal on *ImmPort* and *ImmuneSpace* repositories and may be updated over time. To provide a consistent data source for reproducible results, we also archived a static copy of the data as a “virtual study” in *ImmuneSpace* (Figure 1A & 2).

#### Identification of vaccine study cohorts with transcriptomic profiles

Through a literature search conducted from 2017 to 2020, we identified target publications with systems-level profiling of human vaccination responses. We found 16 HIPC-funded vaccinology studies in *ImmPort* with transcriptomics datasets generated with matching immune response outcomes. Notably, we have supplemented the HIPC data previously available in *ImmPort* by curating and submitting 14 additional human vaccination studies to *ImmPort*. For studies that were not in *ImmPort*/*ImmuneSpace*, we located the underlying data by surveying public transcriptome databases (e.g., Gene Expression Omnibus (GEO)) or reaching out to study authors to request data access, allowing us to submit to *ImmPort* on their behalf. These datasets were then made available via *ImmuneSpace* to be processed for standardization, preprocessing checks, and normalization. The standard analytical pipeline enables reproducibility and comparability of future studies to be correlated with publicly available immune response measurement. This process created the virtual study for the HIPC named the Immune Signatures Data Resource (Figure 1A, Figure 2).

### Gene Expression Data processing pipeline

Data were read directly from *ImmuneSpace* using *ImmuneSpaceR* functions and subsequently preprocessed, quality controlled, and integrated using the following pipeline:

#### Quality Control of Microarray experiments

The *ArrayQualityMetrics* R package^40^ was used for quality control and assurance of all microarray experiments (Figure 3A). Outlier detection was based on the following statistics: i) Mean absolute difference of M-values (log-ratios) of each pair of arrays, ii) the Kolmogorov-Smirnov statistic *K_a_* between each array’s signal intensity distribution and the distribution of the pooled data and, iii) the Hoeffding’s statistic *D_a_* on the joint distribution of A (average) and M values for each array. Using pre-specified criteria within an established public microarray data reuse pipeline^40^, we flagged for removal arrays that failed all three quality control statistics.

**Figure 3:**
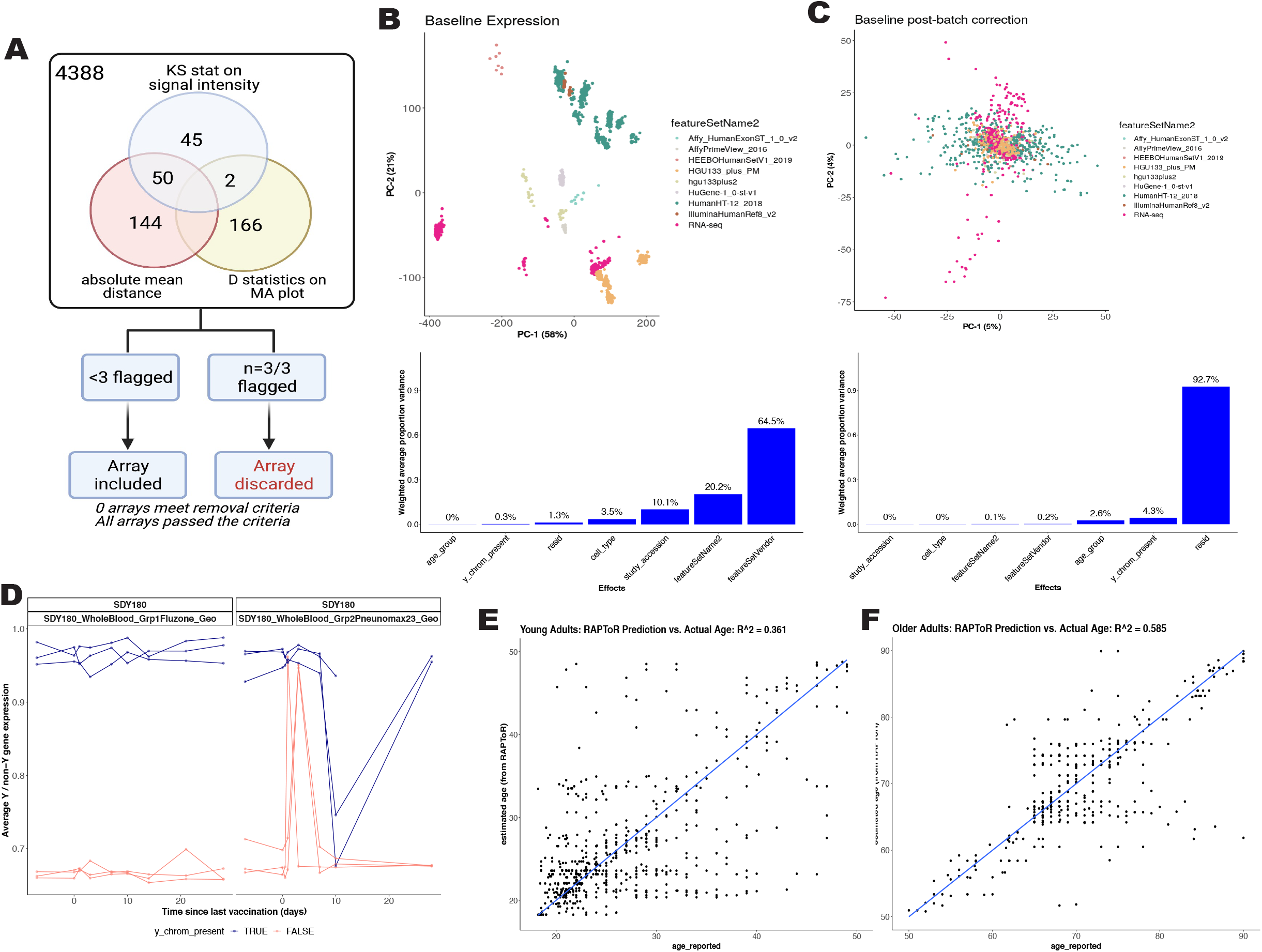
Quality control assessments of transcriptomics data. A. Sample quality assessments of gene expression datasets using Array Quality metrics. Array quality metrics package was employed to assess quality of microarray datasets by checking the following criteria: a.) absolute mean difference between arrays to check the probe and median intensity across all arrays, b.) Kolmogorov-Smirnov statistics to check the signal intensity distribution of arrays, comparing each probe versus distribution of test statistics for all other probes, c.) Hoeffding’s D-statistics for arrays. Arrays were excluded if they fail all three criteria above. B-C: Principal component analysis (Top) and Principal Variation component Analysis (PVCA) of baseline expression data per study before (B) and after batch correction (C). D. Biological sex imputation based on expression of Y-chromosome genes. We used 13 Y-chromosome-associated genes to cluster samples into 2 groups assuming biological male or female. E-F. Age imputation based on transcriptomic profiles for studies without reported ages (SDY1260, SDY1264, SDY1293, SDY1294, SDY1364, SDY1370, SDY1373, SDY984) via the RAPToR R package^45^. Virtual studies were split into young (age < 50, **E**) and older (age >= 50, **F**) for two separate predictive models.

#### Preprocessing

Raw probe intensity data for Affymetrix studies were background-corrected and summarized using the RMA algorithm^41^ while the function *read.ilmn (limma* R package) was used to read and background correct Illumina raw probe intensities. To integrate RNA-seq and microarray data, raw counts for RNA-seq data were converted to log-transformed values incorporating observational level weights to account for technical variations using the *voom*^42^ transformation. Expression data within each study were quantile normalized and log-transformed separately for each cohort/sample type.

#### Annotation

We annotated the manufacturing IDs (probes from microarray/Illumina) to their corresponding gene alias. Gene aliases were mapped to the recent gene symbols from the HUGO Gene Nomenclature Committee^43^ [accessed Dec 23, 2020]. For the rare case where a gene alias mapped to more than one gene symbol, the mapping was resolved by the following: i) If a gene alias mapped to itself as a symbol, as well as other symbols, then it was mapped to itself; ii) if the gene alias mapped to multiple symbols that did not include itself, then the gene alias was dropped from the study. As a result, the raw gene expression matrix was reduced to 10086 HUGO gene aliases with known unique mapping.

#### Gene-based expression profiles

Expression data were summarized at the probe level (for microarray data) and gene-alias level (RNA-seq) to the canonical Gene-Symbol level. The probes / gene-aliases were summarized by selecting the probe or gene-alias with the highest average expression (mean of probes across all samples, take the highest mean) across all samples within the matrix (cohort and sample type).

#### Cross-Study normalization

One of the main assumptions in expression analysis is that differences in gene expression across conditions occur in a relatively small number of processes. As such, the distribution across conditions should be similar, and departures of these assumptions are corrected, for example, using quantile normalization. This procedure usually creates a target distribution using all samples available, but we observed dissimilar distributions in our collection stemming from various platforms used. Such differences lead to extensive distributions and introduce artifacts in the data (Figure 3B and 3C). The target distribution was obtained from samples using Affymetrix platforms, resulting in a well-defined distribution, and each sample in our collection was quantile normalized to this target distribution. Before cross-study normalization, there were 35,725 representative gene symbols present. There were 25,639 genes removed after normalization, as these genes were not present in all the studies. This yielded a final expression matrix of 4795 samples from 1405 participants representing 10,086 genes (Figure 2).

#### Determining and adjusting for technical confounders

We studied the primary sources of variation in the data, including the study effect (which also encompasses the impact of different expression platforms (RNA-seq, Affymetrix arrays, Illumina arrays, etc.), sample types (Whole blood, PBMC), as well as demographics. We conducted Principal Component Analysis (PCA) to visualize such associations in a bidimensional space of principal components (PCs) and applied Principal Variance Component Analysis (PVCA)^44^ to quantify the amount of variability attributed to different experimental conditions. This approach models the multivariate distribution of the PCs computed for the PCA as a function of experimental factors and estimates the total variance explained by each factor via mixed-effect models. Since many studies included only one vaccine, temporal variations due to vaccine response were confounded with the study effect. The assessment of the primary technical sources of variation was carried out using only the pre-vaccination data, not affected by the targeted pathogen and vaccine type used in the different studies. Of note, all studies enrolled healthy volunteers, and the first biosample was obtained pre-vaccination. The targeted pathogen and vaccine type should not affect these baseline data. Platform, study, and sample types were identified as significant sources of variation in the gene expression matrix. The effect of those three variables was estimated by modeling gene expression at baseline (at which no vaccine or timepoint effect exists) with a linear model using the *limma* framework, including feature set vendor (Platform/Affy), study (batch factors), and sample type, Y-chromosome genes presence, as covariates. Study and cell-type effects were estimated using a linear model with age, Y-chromosome genes presence (biological sex), study, sample type (Whole Blood/PBMC), study, and platform as additive effects. From here, the study, platform, and cell-type effects were eliminated from the entirety of the expression matrix. There were three studies (SDY1276, SDY1264, SDY180) that contained multiple cohorts and were treated as separate studies.

#### Biological sex imputation

Imputation of biological sex, as defined by the presence of a Y-chromosome, was carried out based on the gene expression profiles of 13 Y-chromosome genes. Within each study, a multidimensional scaling was first applied to the Y-chromosome gene expression profiles. K-means clustering was then used to cluster samples into two groups. Participants in the cluster with higher mean expression values were considered male (i.e., the Y-chromosome was present) while those in the cluster with lower expression were considered female (i.e., the Y-chromosome was absent). The consistency of the Y-chromosome presence assignment across time points was verified (Figure 3D). In the (few) cases where imputation was not in agreement across all time points, the reported sex was used and if no sex was reported, imputation followed a majority rule principle.

#### Age Imputation

Age imputation for studies without reported ages (SDY1260, SDY1264, SDY1293, SDY1294, SDY1364, SDY1370, SDY1373, SDY984) employed the RAPToR R v1.1.5 package^45^. The RAPToR algorithm takes in a reference set of gene expression time series with reported ages and generates a near-continuous, high-temporal resolution from the interpolated reference dataset. Transcriptomic profiles of participants without reported ages were compared to the reference dataset via a correlation profile, providing age estimates for the sample. Finally, random subsets of genes from the subject’s transcriptomic profile were bootstrapped to ascertain a confidence interval for the imputed age. We generated the reference dataset using the transcriptomic profiles of 21 studies in our resource for which age was reported. The studies were split into younger (age < 50) and older (age ≥ 50) cohorts, thus two different models were generated, and only baseline transcriptomic profiles were used in the reference dataset. As RAPToR also enables phenotypic data to be incorporated into the interpolation model, each possible combination of phenotypic features was tested. For each combination, RAPToR predicted the age for participants in the 21 studies with known age, and the goodness of fit was evaluated by the coefficient of determination (R^2^). The best model for the younger and older cohorts was then used to impute ages for the 7 studies without reported age (Figure 3E, 3F)

### Immune response datasets processing pipeline

To identify the molecular signatures that correlate with vaccine immunogenicity, we included immune response readouts in the creation of this data resource. For studies that were missing vaccine response endpoints in their public data deposition, we contacted study authors and requested available antibody response measures to vaccine antigens. Once shared, these data were submitted to *ImmPort* and linked to the relevant studies. These readouts include neutralizing antibody titers (Nab), hemagglutination inhibition assay (HAI) results for influenza studies, and Immunoglobulin IgG ELISA assay results. In participants for whom the humoral immune response was measured with multiple assays, the preference was given to HAI for influenza or Nab for non-influenza studies, then IgG ELISA datasets. The antibody measures were normalized within each study by estimating the fold-change differences between the post-vaccination time-point (generally between day 28 or day 30) compared to the baseline measurement. For influenza studies where the vaccine included multiple strains, the fold changes between the post-vaccination versus baseline were calculated for each strain, and the maximum fold change (MFC) over the strains was selected^34^. Due to the variability in baseline antibody (Ab) levels and immune memory such as influenza vaccines, we also estimated the maximum residual after baseline adjustment (maxRBA) method by calculating the maximum residual across all vaccine strains to adjust for variable baseline Ab levels using the R package *titer*^21^. A total of 30 studies with 1405 participants and 4795 samples have both transcriptomics and immune response readout data available (Figure 2). This dataset enables researchers to carry out comparative analyses using immunogenicity data as well as prediction of the quality of response across multiple vaccines.

### Data Records

The Immune Signatures Data Resource is available online for download by the research community from this website: The data is hosted on *ImmuneSpace* and can be accessed via the R package *ImmuneSpaceR* (https://rglab.github.io/ImmuneSpaceR/). The resource is available for use by the scientific community and can be downloaded from a research data repository IS2 https://www.ImmuneSpace.org/is2.url. A summary of datasets, with their corresponding study ID and accession numbers, is provided in Table 4.

### Technical Validation

#### Quality Control and Assurance

For global quality control across all public microarray data, we used a well-established pipeline available through the *ArrayQualitymetrics* R package^40^. Using pre-specified criteria established in the existing public microarray data reuse pipeline^46^, arrays that failed 3 out of 3 calculated quality control statistics were flagged for removal (see Methods). Consistent with standard practice to perform such quality control analysis prior to downstream analysis and dataset submission to the Gene Expression Omnibus, none of the samples were outliers by all three statistics (Fig 3A). As expected for data from published peer-reviewed studies, all the identified studies passed the quality assurance method using the *Arrayqualitymetrics* method.

#### Y-chromosomal presence and age imputation

A few studies were missing information for sex and for age. To achieve data completeness, we included the biological sex imputation based on the imputed presence of the Y-chromosome using gene expression, as well as imputation of age when the variable was missing or defined by a broad range of values. Age imputation employed the RAPToR tool using 21 studies with reported age to define the best predictive model for the younger (age < 50 years) and older (age ≥ 50 years) cohorts separately. The highest correlation coefficients from the young cohort were generated by taking into account the model (X ~ age_reported + matrix) with a correlation coefficient of R^2^=0.367 (Figure 3E), while the old cohort yielded a prediction R^2^ of 0.536 for their highest coefficient value (Figure 3F).

#### Definition of Vaccination Studies Transcriptomic Cohort

Data preprocessing in *ImmuneSpace* yielded a total of 30 studies and 59 cohorts, with 1482 participants and 5413 samples. After the data was preprocessed and quality control measures were performed, we further assessed the identified cohorts as defined in the flow diagram (Figure 2). This curation included: i) removing participants that were not relevant to the objective (n=34); ii) removing samples due to inconsistencies with time design determination (n= 178); iii) removing participants with no baseline expression data (n=42). Some studies, such as SDY1368 and SDY67, were dropped from the normalized data sets as they did not include subjects within our target age range (18-50 years). In summary, we report that the final Immune Signatures Data Resource contains 53 cohorts from 30 studies with 1405 participants and 4795 samples.

#### Assessment and adjustment of the batch effects

We evaluated the main sources of variation on the gene expression matrix to identify and adjust technical confounders (RNA-seq, Affymetrix arrays, Illumina arrays, etc.), study, and specimen types (e.g., whole blood vs. PBMCs) using the baseline samples. Since all studies enrolled healthy volunteers, and the first sample was taken pre-vaccination, pathogen and vaccine type would not affect the baseline data. Figure 3B clearly demonstrates robust clustering of samples by study, which are also grouped by platform type. The study effect and type of platform used accounted for the vast majority (95%) of variation, followed by specimen types (3.6%). It is thus essential that the data are corrected for these major effects prior to any analytical usage [see Materials and Methods for further details]. The study, platform type, and specimen type-specific effects were estimated using a linear model that also included age and Y-chromosome presence as additive effects using only baseline expression. Once the study, platform, and specimen-type effects were estimated, they were eliminated from the entirety of the expression matrix. Figure 3B shows that those effects can successfully be adjusted from the data, thus leading to a matrix of expression that is free of most technical biases induced by the laboratory and cell-type effects.

#### Immune Signatures Transcriptomics and Immune Response Datasets

We report the total number of assay samples collected from the transcriptomic and immune response datasets tallied by targeted pathogen and vaccine type, across multiple systems vaccinology datasets (Figure 4A). We captured about ~3000 HAI antibody titer results from influenza studies that were measured by the standard HAI assay pre- and at multiple time points post-vaccination, depending on the study. Mean titers were calculated for the reported strains of the virus and were based on the highest dilution reported at day 28-30 post-vaccination. In addition, neutralizing antibody (NAB) titers and IgG ELISA results specific to each pathogen were determined by each study and are summarized (Figure 4A). The overall transcriptomics dataset comprises multiple time points from 7 days pre-vaccination up to day 180 days post-vaccination (Figure 4B). While most of the datasets focus on the young adult population (ages 18-50 years old), the data resource also includes studies that profile older adults following hepatitis B, influenza, and varicella vaccination (Figure 4C) that may be useful for analysis. The Euler diagram describes the dataset overlap of participants with transcriptomics datasets and corresponding to one or more immune response datasets (Figure 4D).

**Figure 4:**
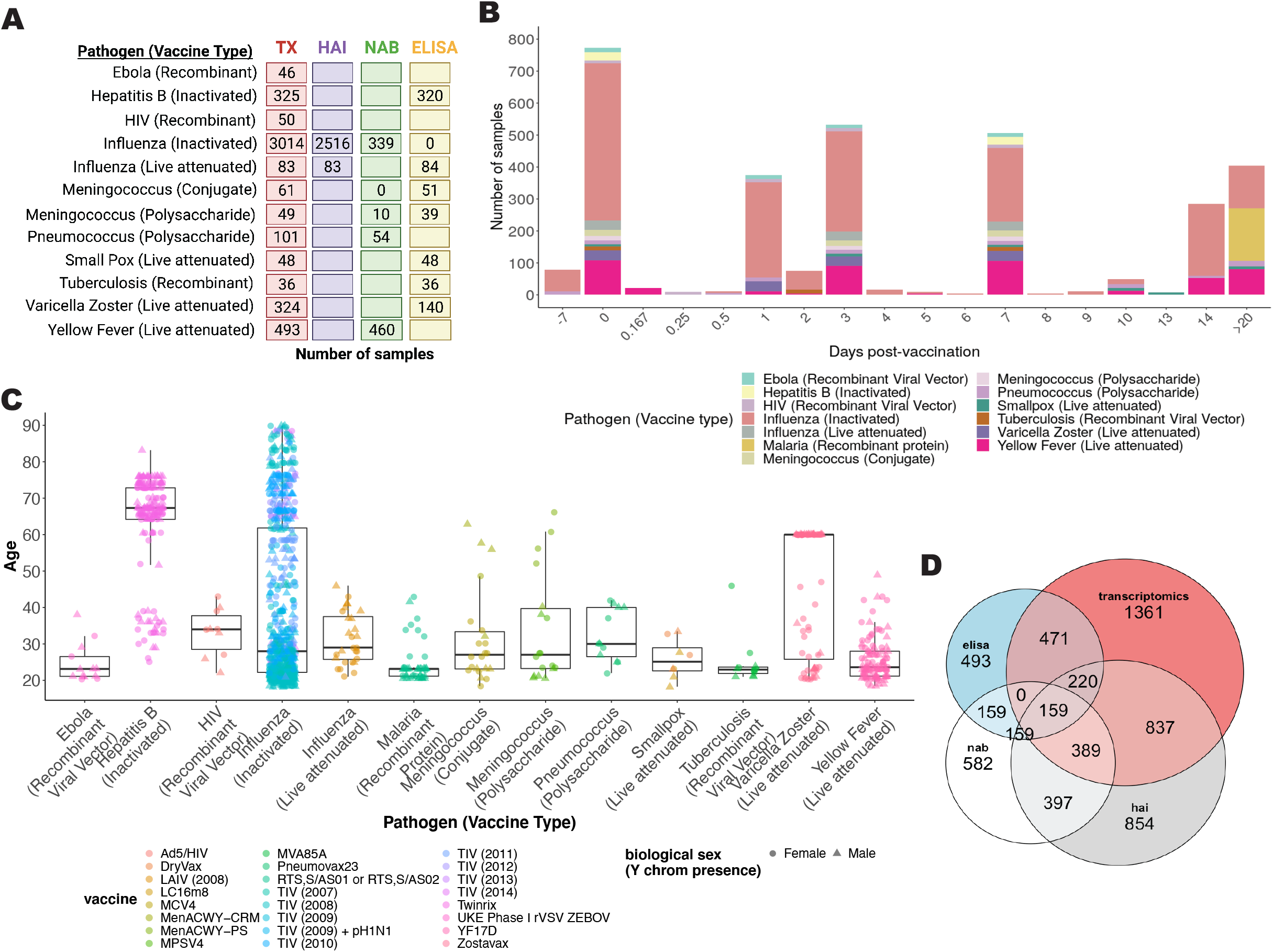
Immune Signatures Transcriptomics Overview for young and old datasets. A. Number of samples available for each data type, including transcriptomics (TX), hemagglutination inhibition assay (HAI), neutralizing antibody assay (NAB), and ELISA assays (ELISA). B. Bar plot depicting the number of samples at each time point. The colors within each bar indicate the breakdown for each unique combination of pathogen and vaccine type. Day −7 and day 0 correspond to times pre-vaccination. C. Box plot depicting the participant’s age distribution for each unique combination of pathogen and vaccine type D. Each area-proportional Euler diagram represents the total number of participants with corresponding data types.

Heterogeneity of the immune response to vaccination across targeted pathogens and vaccine types was reflected in variation in the longitudinal trajectories of HAI and NAB titer measurements (Figure 5A and 5B). HAI and NAB titers generally increased by 14-28 days after vaccination but attenuated at different times for each vaccine (Figure 5A and 5B). Change in NAB titers after vaccination were significantly different across the 5 unique combinations of targeted pathogen and vaccine types where these measurements were reported (ANOVA p <10^-10^), with significant differences across all 5 groups except between meningococcus and yellow fever vaccines (Figure 5C). Some influenza vaccination studies reported both HAI and NAB measures of immunogenicity, and there was a significant positive correlation between the vaccination-induced changes in these titers across participants (Spearman’s rho = 0.45, p <10^-10^) (Figure 5D).

**Figure 5:**
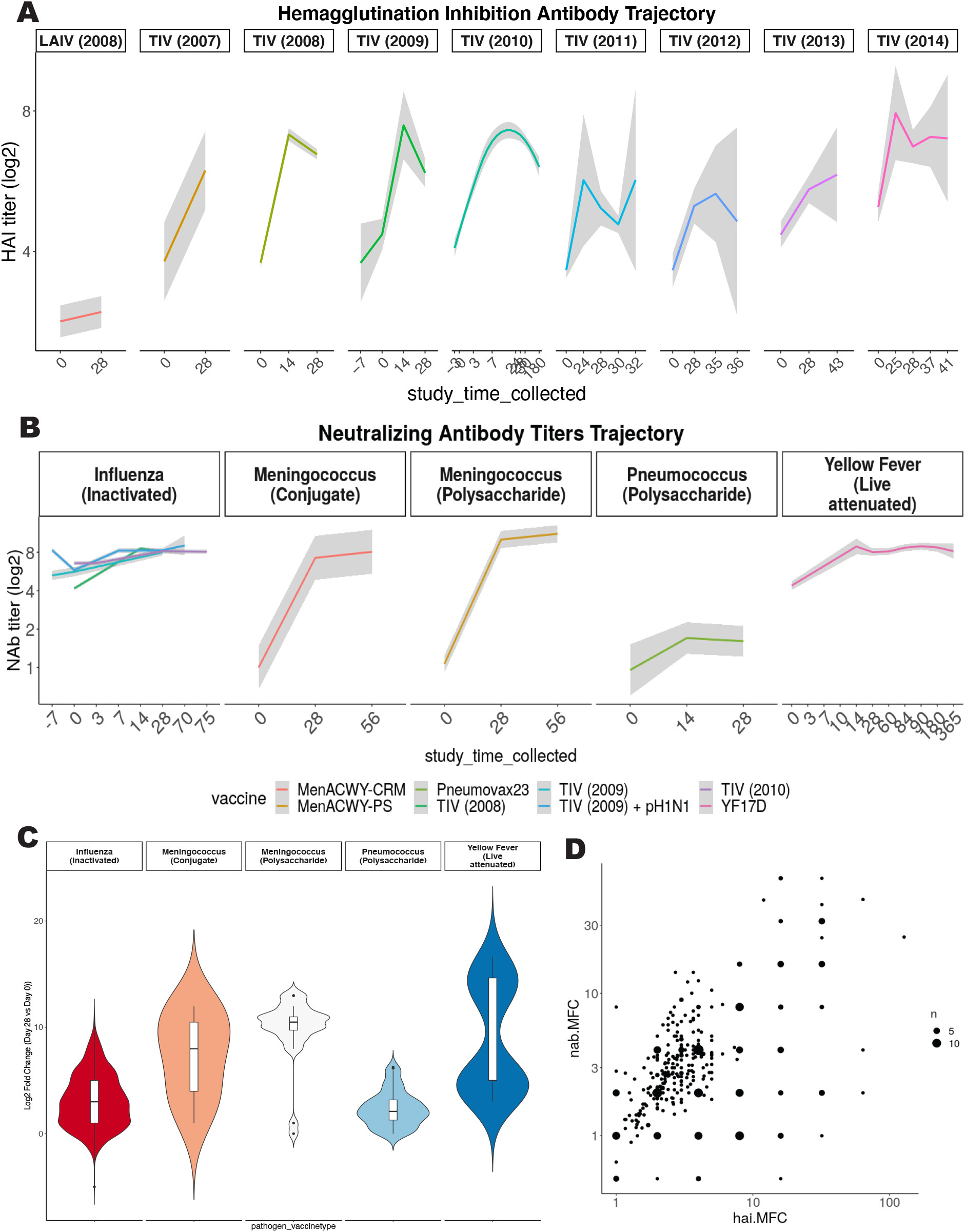
Immune Response Dataset Overview. A. The longitudinal trajectory (summarized as a loess curve) of hemagglutinin inhibition assay (HAI) measurements (in log_2_ scale) by influenza vaccine type and year. B. The longitudinal trajectory of neutralizing antibody (NAB) titers (in log2 scale) for influenza, meningococcus, pneumococcus, and yellow fever vaccines. C. Neutralizing antibody titers were plotted for each unique combination of targeted pathogen and vaccine type to compare each participants’ post-vaccination (day 28-30) values versus baseline (day 0). The violin plot shows the variation in magnitude for each unique combination of targeted pathogen and vaccine type. D. The correlation plot of influenza studies compares the maximum fold change (MFC) across strains for hemagglutinin inhibition assay (HAI) titers versus neutralizing antibody (NAB) titers. Size is proportional to the number of samples analyzed.

#### Usage Notes

The expression data and accompanying meta-data have been made available with different formats and options to ease usage. Data are available as standard expression sets (eSet) objects, the R/Bioconductor structure unifying expression values, metadata, and gene annotation. Both normalized data and batch-adjusted data are available (Table 4). Users interested in a single study or those planning to work exclusively within participants’ changes may opt for the normalized data without batch adjustment. For comparison of time points across studies or developing algorithms that use expression data, batch corrected matrices should be employed. Imputed age values for participants with no reported age were included to facilitate the use of age as a covariate in future analysis. Such analysis can be carried out with the complete data set and can be followed up by a sensitivity analysis using the small cohort with age-reported data. For the use of expression sets with the corresponding immune response per participant, these are available in eSets noted with a response. The selected immune response outcome per study is also summarized in Table 3.

**Table 1:**
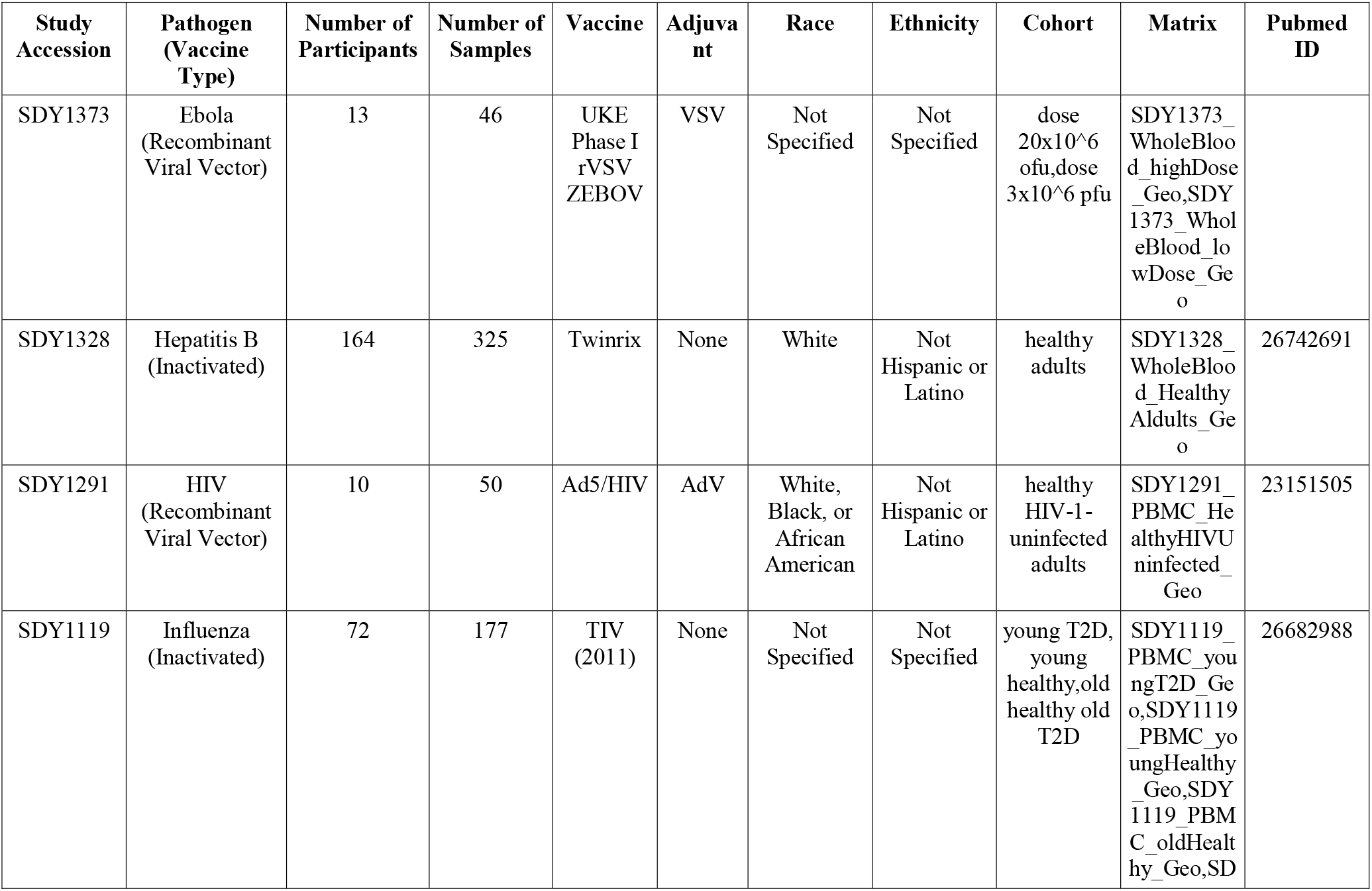
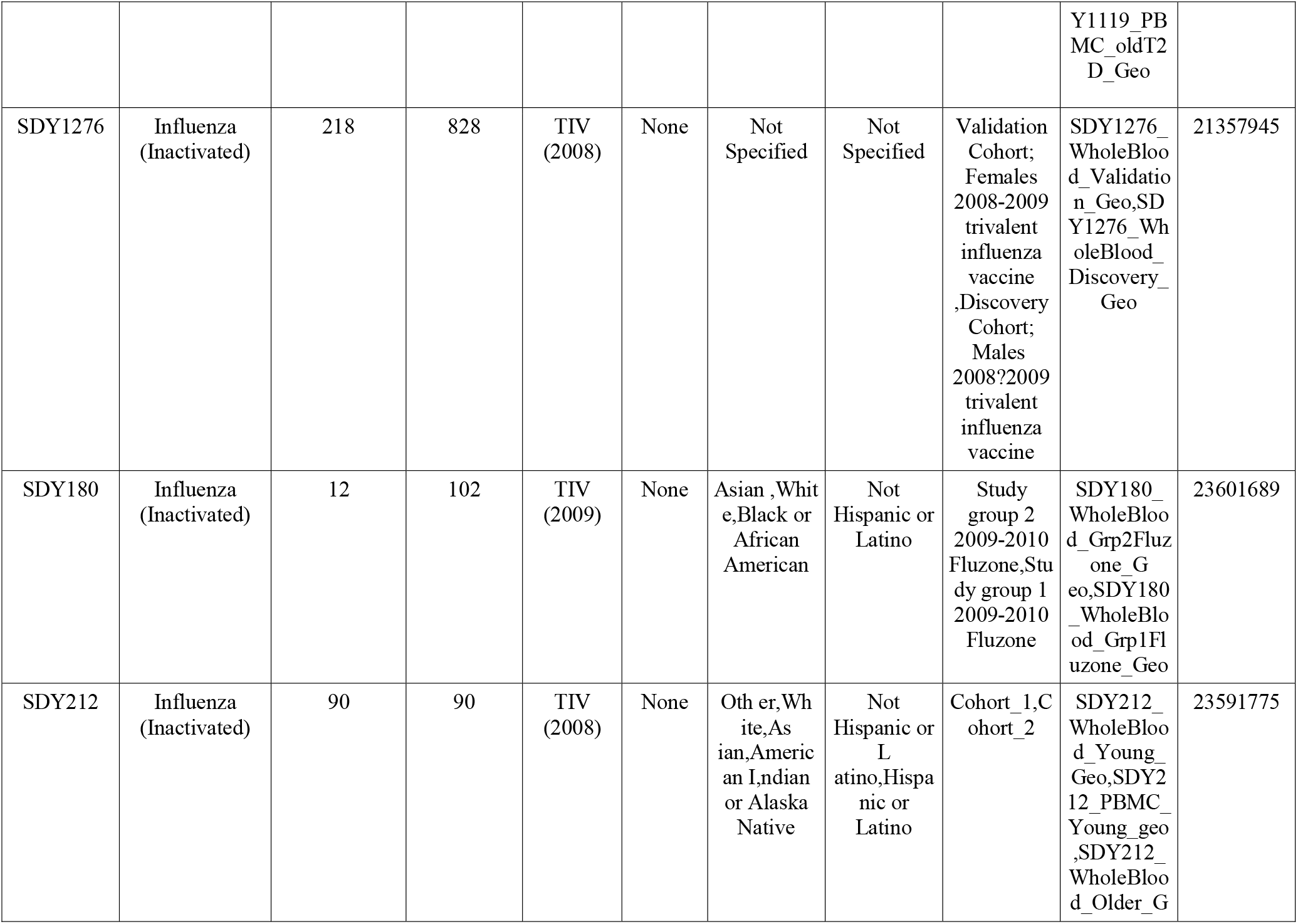
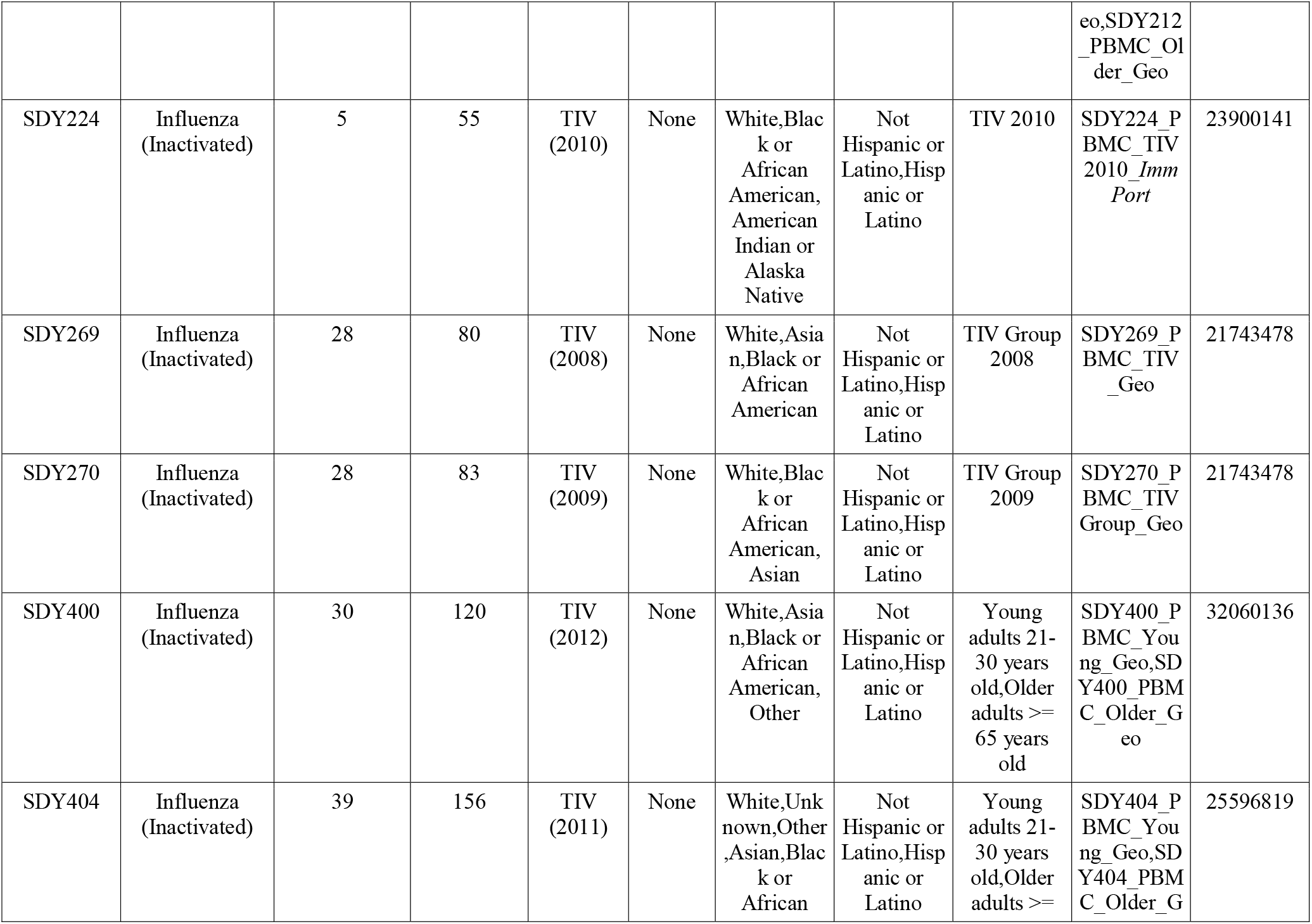
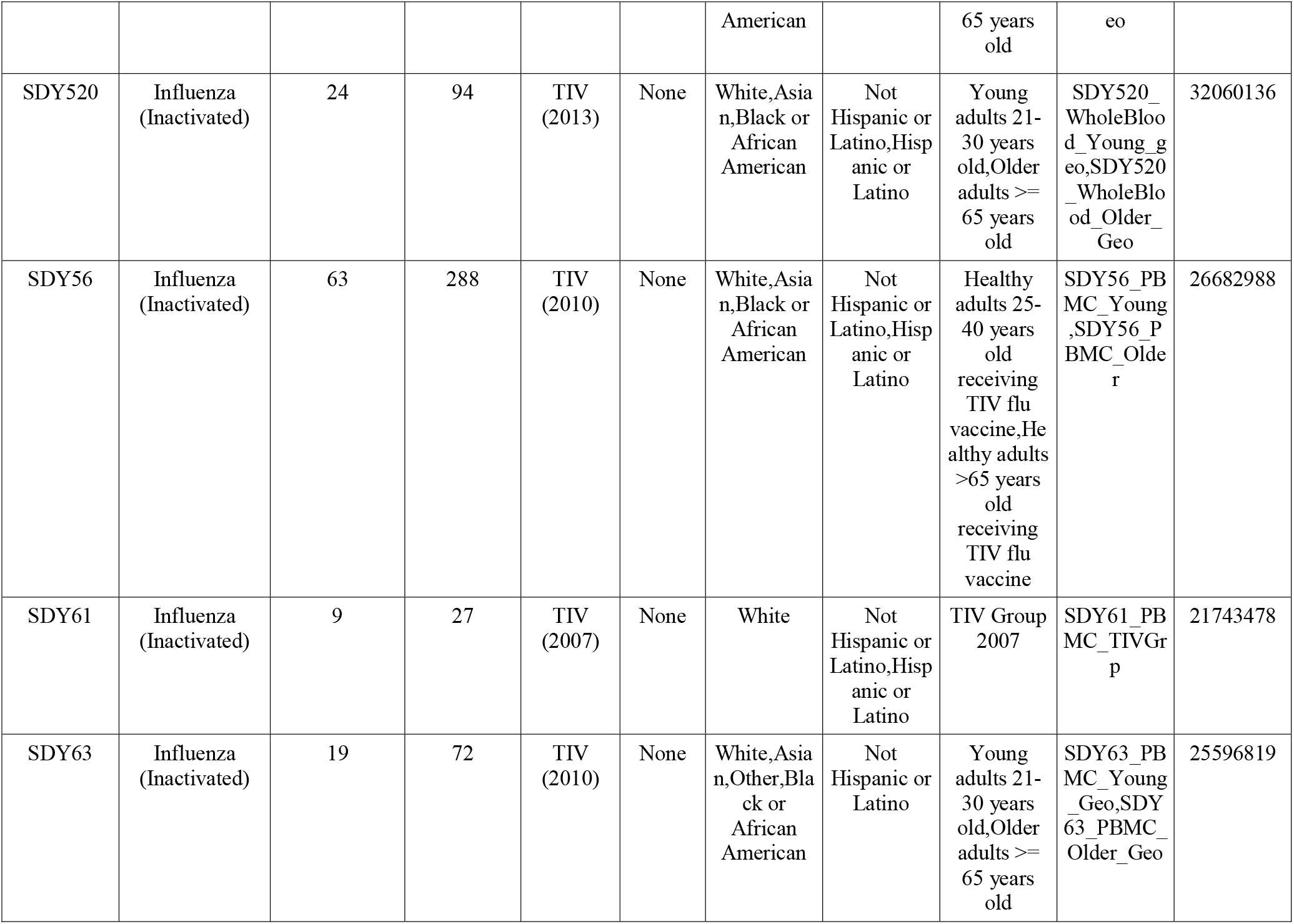
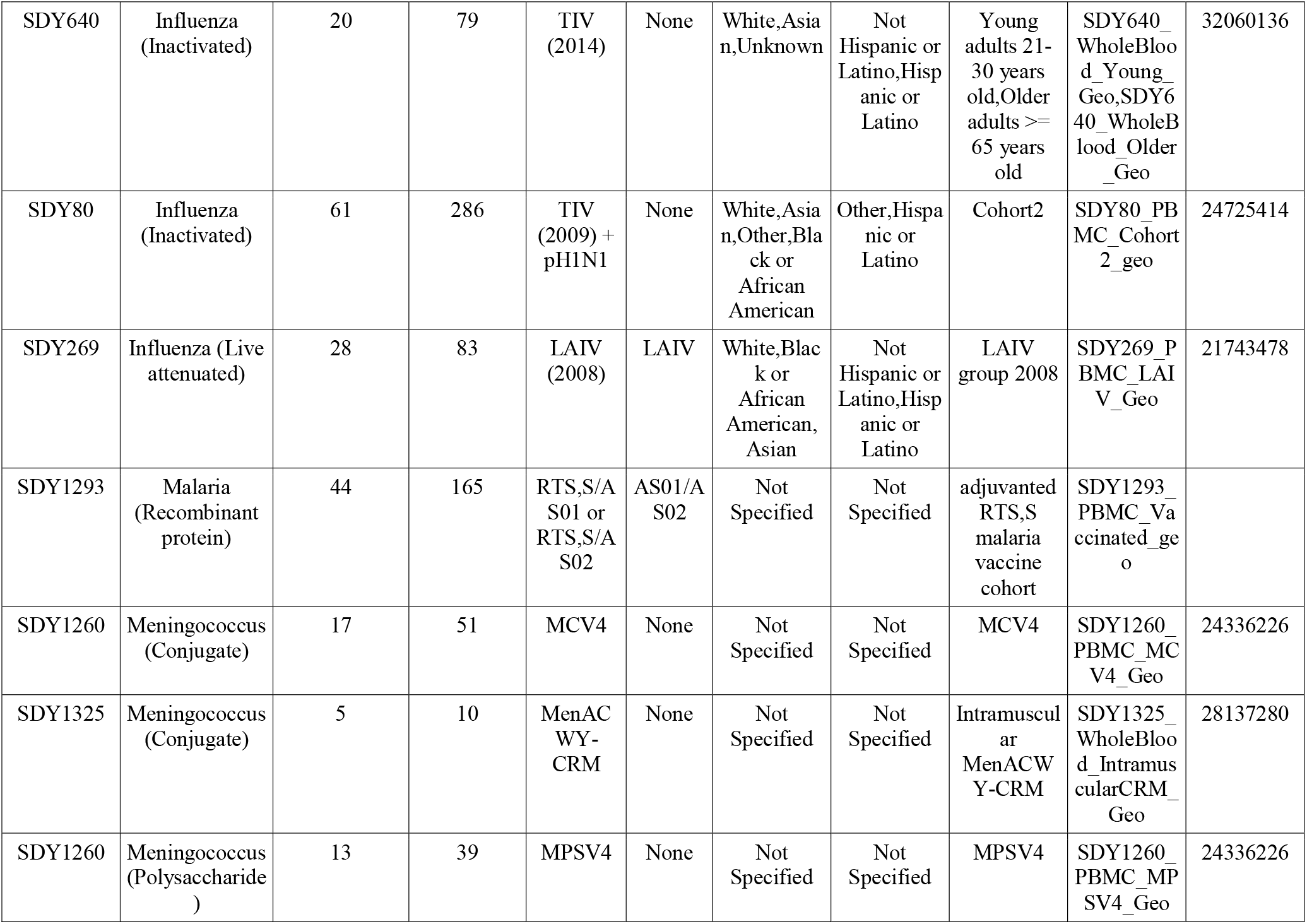
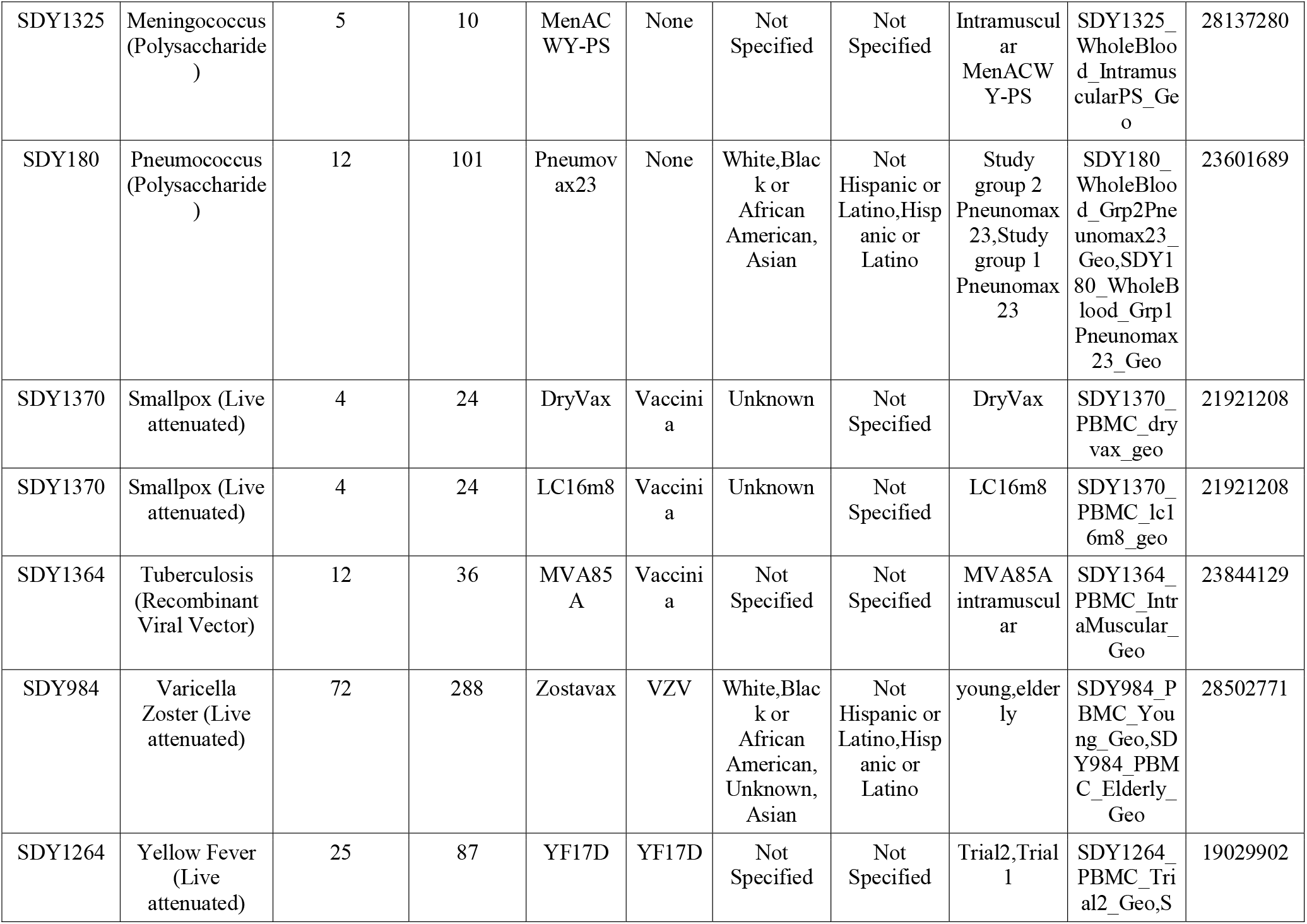
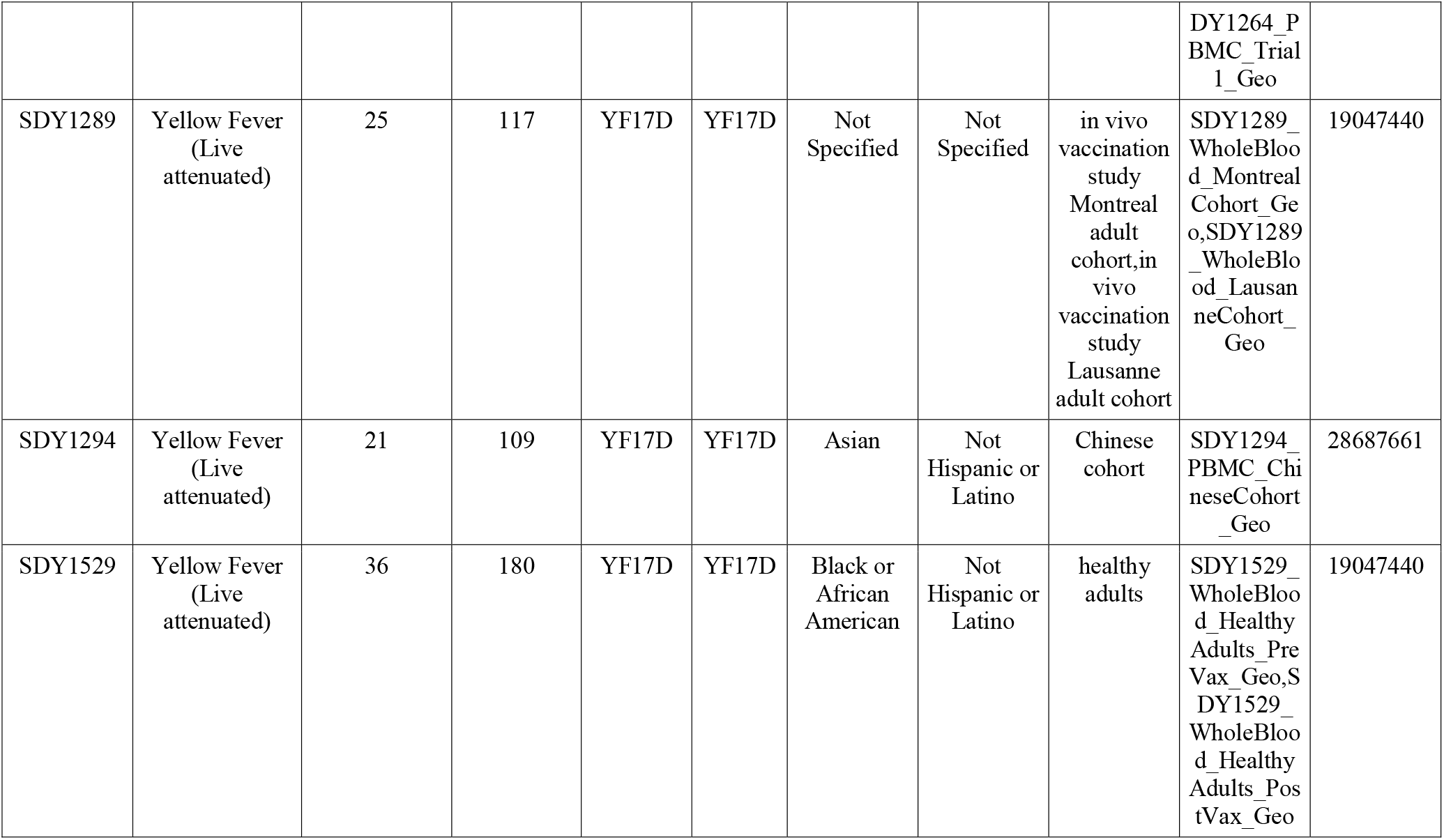
Overview of Immune Signatures Data Resource Study Participants Metadata.

**Table 2:**
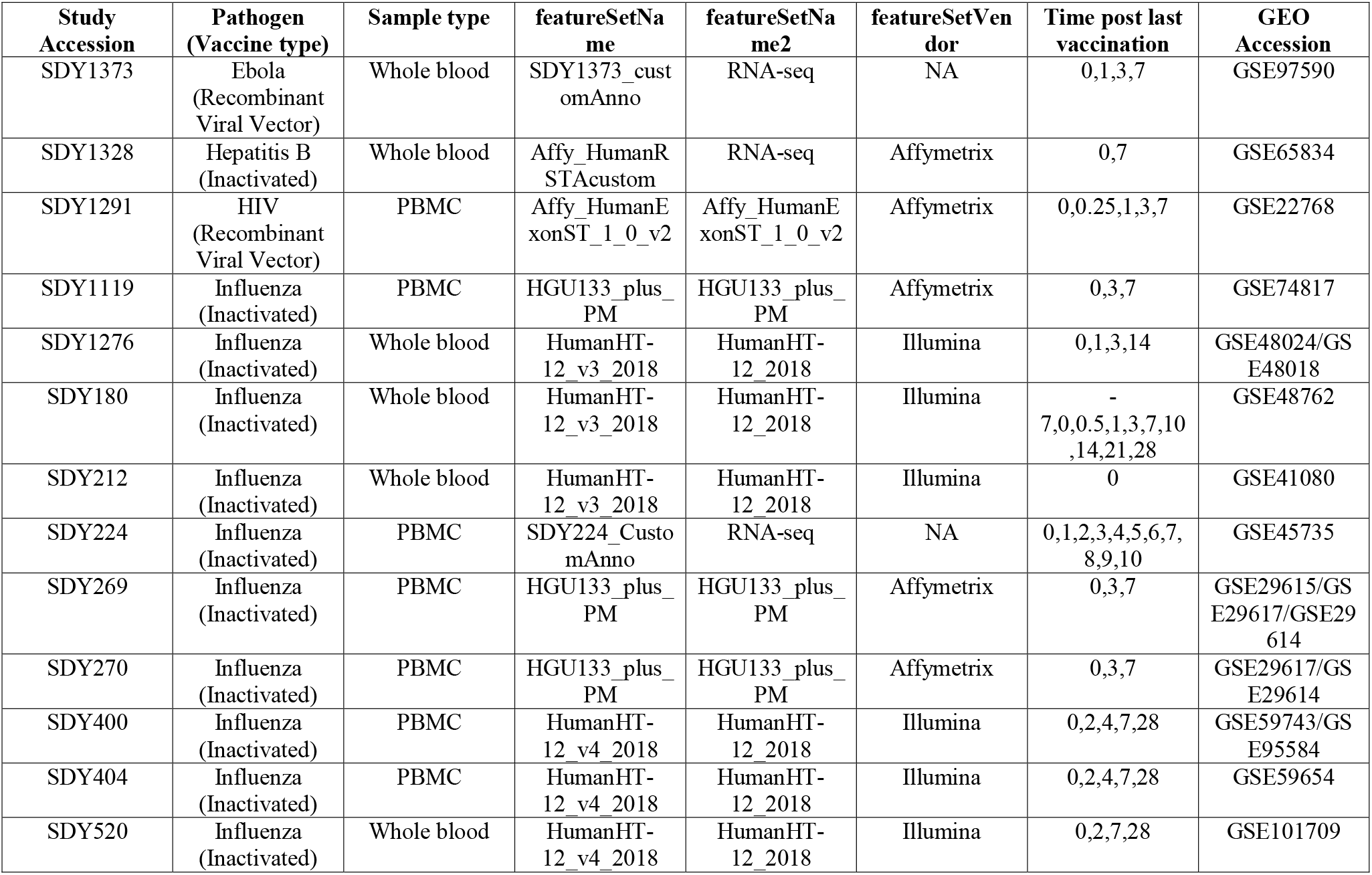
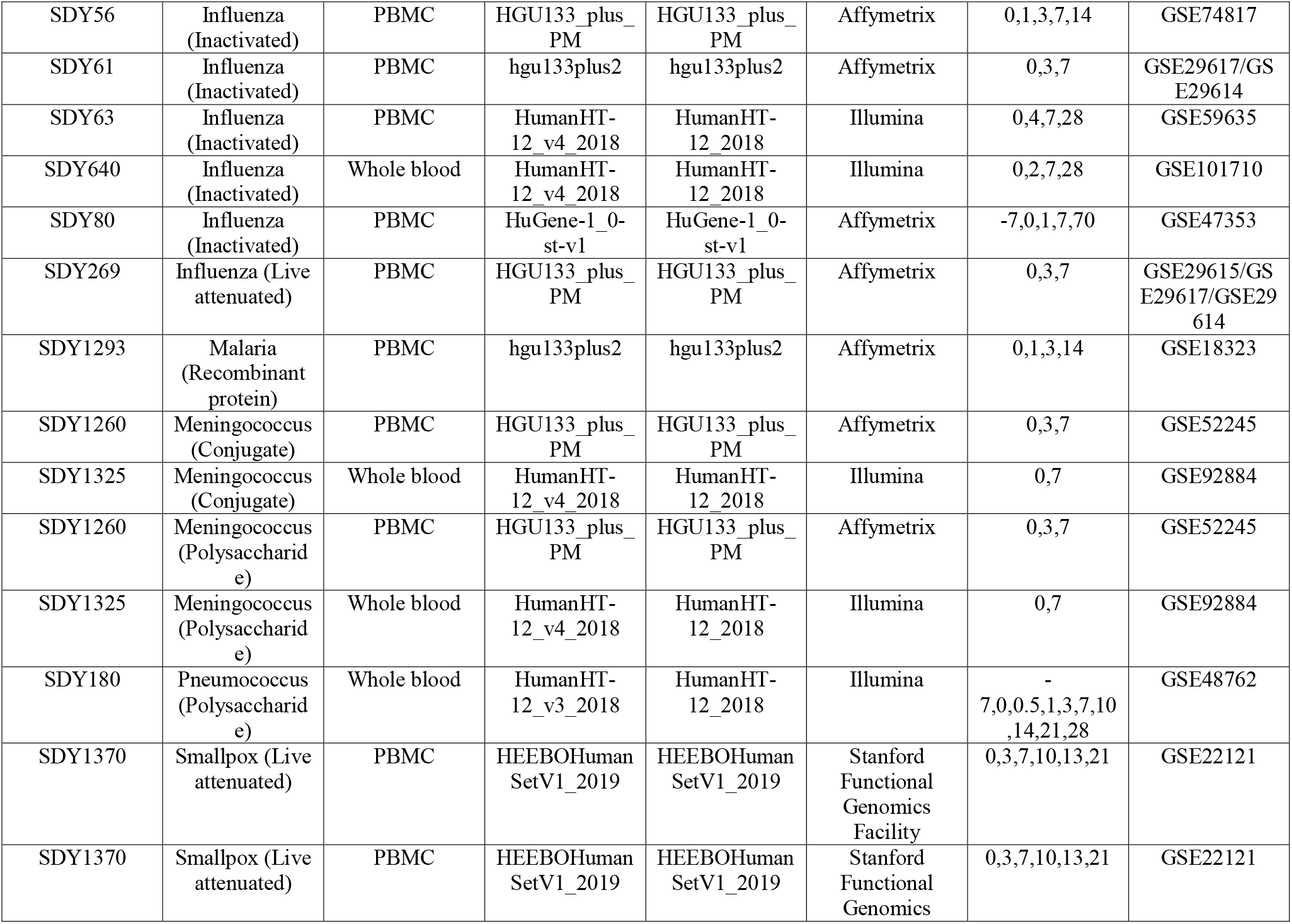
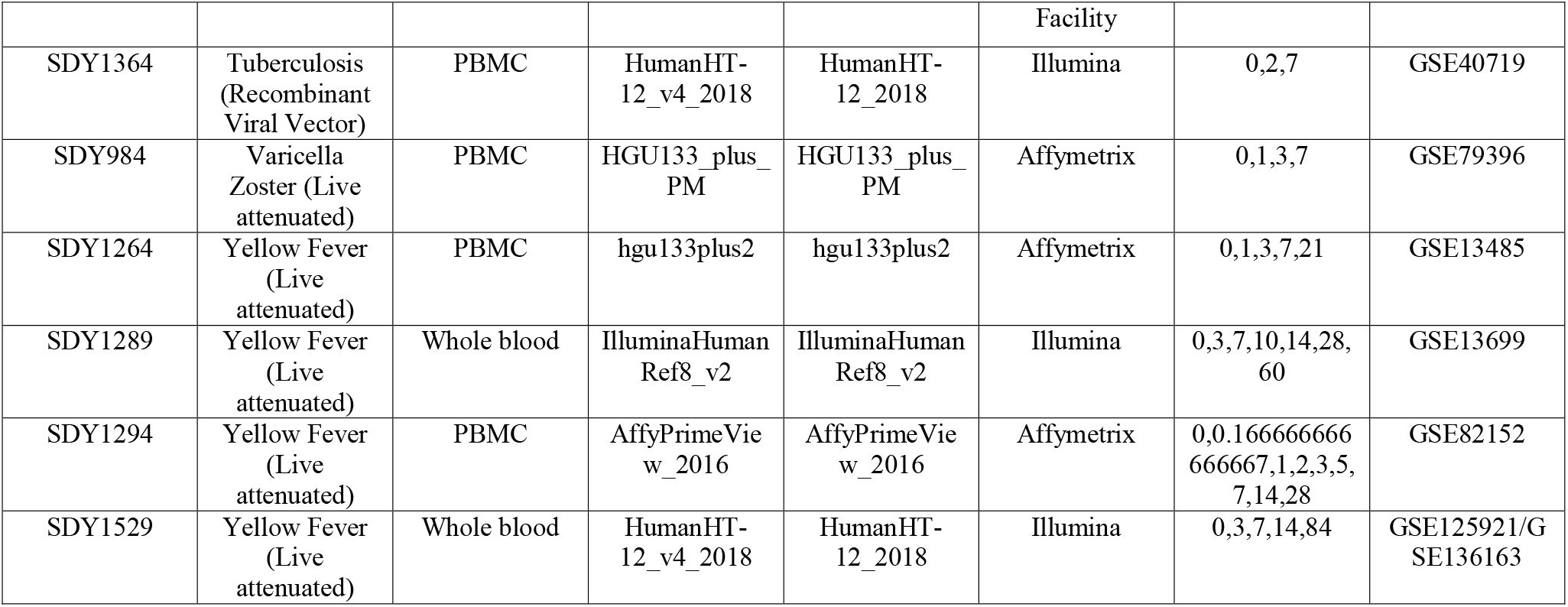
Overview of Transcriptomics Datasets Included in the Resource.

**Table 3:** Studies with corresponding Immune Response Data.

| **Study Accession** | **Pathogen Vaccine type** | **Number of Participants** | **Number of Samples** | **Selected Immune Response Assay** |
| --- | --- | --- | --- | --- |
| SDY1328 | Hepatitis B (Inactivated) | 160 | 320 | ELISA |
| SDY1119 | Influenza (Inactivated) | 72 | 177 | HAI |
| SDY1276 | Influenza (Inactivated) | 214 | 816 | HAI, NAb |
| SDY180 | Influenza (Inactivated) | 12 | 102 | HAI, NAb |
| SDY212 | Influenza (Inactivated) | 88 | 88 | HAI |
| SDY224 | Influenza (Inactivated) | 5 | 55 | HAI |
| SDY269 | Influenza (Inactivated) | 28 | 80 | HAI |
| SDY270 | Influenza (Inactivated) | 28 | 83 | HAI |
| SDY400 | Influenza (Inactivated) | 30 | 120 | HAI |
| SDY404 | Influenza (Inactivated) | 39 | 156 | HAI |
| SDY520 | Influenza (Inactivated) | 24 | 94 | HAI |
| SDY56 | Influenza (Inactivated) | 30 | 148 | HAI |
| SDY61 | Influenza (Inactivated) | 9 | 27 | HAI |
| SDY63 | Influenza (Inactivated) | 19 | 72 | HAI |
| SDY640 | Influenza (Inactivated) | 20 | 79 | HAI |
| SDY67 | Influenza (Inactivated) | 159 | 477 | HAI |
| SDY80 | Influenza (Inactivated) | 60 | 281 | NAb |
| SDY269 | Influenza (Live attenuated) | 28 | 83 | HAI |
| SDY1260 | Meningococcus (Conjugate) | 17 | 51 | ELISA |
| SDY1325 | Meningococcus (Conjugate) | 4 | 8 | NAb |
| SDY1260 | Meningococcus (Polysaccharide) | 13 | 39 | ELISA |
| SDY1325 | Meningococcus (Polysaccharide) | 5 | 10 | NAb |
| SDY180 | Pneumococcus (Polysaccharide) | 6 | 54 | NAb |
| SDY1370 | Smallpox (Live attenuated) | 4 | 24 | ELISA |
| SDY1370 | Smallpox (Live attenuated) | 4 | 24 | ELISA |
| SDY1364 | Tuberculosis (Recombinant Viral Vector) | 12 | 36 | ELISA |
| SDY984 | Varicella Zoster (Live attenuated) | 35 | 140 | ELISA |
| SDY1264 | Yellow Fever (Live attenuated) | 25 | 87 | NAb |
| SDY1289 | Yellow Fever (Live attenuated) | 14 | 84 | NAb |
| SDY1294 | Yellow Fever (Live attenuated) | 21 | 109 | NAb |
| SDY1529 | Yellow Fever (Live attenuated) | 36 | 180 | NAb |

**Table 4:** List of data files for the Immune Signatures Data Resource.

| File name | Description |
| --- | --- |
| all_noNorm_eset.rds | Gene expression matrix of all participants, log2-normalized expression |
| all_noNorm_withResponse_eset.rds | Gene expression matrix of all participants with matched immune response data, log2-normalized expression |
| all_norm_eset.rds | Gene expression matrix of all participants that are cross-study normalized and batch corrected |
| all_norm_withResponse_eset.rds | Gene expression matrix of all participants with matched immune response dataset, cross-study normalized and batch corrected |
| young_noNorm_eset.rds | Gene expression matrix of participants aged 18-50, log2-normalized |
| young_noNorm_withResponse_eset.rds | Gene expression matrix of participants aged 18-50 with matched immune response data, log2-normalized |
| young_norm_eset.rds | Gene expression matrix of participants aged 18-50, cross-study normalized and batch corrected |
| young_norm_withResponse_eset.rds | Gene expression matrix of participants aged 18-50 with matched immune response data, cross-study normalized and batch corrected |
| old_noNorm_eset.rds | Gene expression matrix of participants aged 60-90, log2-normalized |
| old_noNorm_withResponse_eset.rds | Gene expression matrix of participants aged 60-90 with matched immune response data, log2-normalized expression |
| old_norm_batchCorrectedFromYoung_eset.rds | Gene expression matrix of participants aged 60-90, cross-study normalized and batch corrected using age correction coefficients from young |
| old_norm_batchCorrectedFromYoung_withResponse_eset.rds | Gene expression matrix of participants aged 60-90 with matched immune response data, cross-study normalized and batch corrected using age correction coefficients from young |
| extendedOld_noNorm_eset.rds | Gene expression matrix of participants aged 50-90, log2-normalized expression |
| extendedOld_noNorm_withResponse_eset.rds | Gene expression matrix of participants aged 50-90 with matched immune response data, log2-normalized counts |
| extendedOld_norm_batchCorrectedFromYoung_eset.rds | Gene expression matrix of participants aged 50-90, log2-normalized expression |
| extendedOld_norm_batchCorrectedFromYoung_withResponse_eset.rds | Gene expression matrix of participants aged 50-90 with immune response data, cross-study normalized, and batch corrected using correction coefficients from young |

## Code Availability

The source codes for the Immune Signatures Data Resource and all data are available in ImmuneSpace (https://www.immunespace.org/is2.url). Pre-processing code and supplementary data can be found in the ImmuneSignatures2 R package hosted on Github (https://github.com/RGLab/ImmuneSignatures2).

## Acknowledgments

This research was conducted within the Human Immunology Project Consortium (HIPC) and supported by the National Institute of Allergy and Infectious Diseases. This work was supported in part by NIH grants U19AI128949, U19AI118608, U19AI118626, and U19AI089992, U19AI090023, U19AI089992, U19AI128914, U19AI118610, U19AI128913. The HIPC projects are listed at https://www.immuneprofiling.org/hipc/page/showPage?pg=projects. This work was supported in part by the Canadian Institutes of Health Research [funding reference number FDN-154287]

## Contributions

All authors identified the datasets, performed quality control, and assurance and analyzed the datasets. JD-A, HM, SHK and MSF led the writing and organization of the manuscript. HM, EH, PD, together with the *ImmuneSpace* and *ImmPort* team, implemented the pipeline for data access and visualization. The HIPC Consortium contributed to the conception and design of the work, as well as the acquisition of data. A full list of the HIPC Consortium members can be found in Supplementary File 1. All authors edited and approved the manuscript.

## Ethics declaration

S.H.K. receives consulting fees from Northrop Grumman and Peraton. OL is an inventor on several patents relating to vaccine adjuvants and human *in vitro* systems predicting vaccine action. R.G. has received consulting income from Illumina, Takeda, and declares ownership in Ozette Technologies and Modulus Therapeutics. The other authors declare no competing interests.

## Notes

https://www.immunespace.org/is2.url

