## Supplementary File 1 for "The Immune Signatures Data Resource: A compendium of systems vaccinology datasets"

**Human Immunology Project Consortium (HIPC)**

^&^Members of the HIPC Steering Committee: Alison Deckhut-Augustine (NIAID, NIH, Bethesda, MD, USA); Raphael Gottardo (Fred Hutchinson Cancer Research Center, Seattle, WA, USA); Elias K. Haddad (Drexel University, Philadelphia, PA, USA); David A. Hafler (Yale School of Medicine, New Haven, CT, USA); Eva Harris (University of California, Berkeley, Berkeley, CA, USA); Donna Farber (Columbia University Medical Center, New York, NY, USA); Steven H. Kleinstein (Yale School of Medicine, New Haven, CT, USA); Ofer Levy (Boston Children's Hospital, Harvard Medical School, Boston, MA, USA); Julie McElrath (Fred Hutchinson Cancer Research Center, Seattle, WA, USA); Ruth R. Montgomery (Yale School of Medicine, New Haven, CT, USA); Bjoern Peters (La Jolla Institute for Immunology, La Jolla, CA, USA.); Bali Pulendran (Stanford University School of Medicine, Stanford University, Stanford, CA, USA); Adeeb Rahman (Icahn School of Medicine at Mount Sinai, New York, New York, USA); Elaine F. Reed (David Geffen School of Medicine at University of California, Los Angeles, CA, USA); Nadine Rouphael (Emory University School of Medicine, Atlanta, GA, USA); Minnie Sarwal (University of California, San Francisco, San Francisco, CA, USA); Rafick Sekaly (Emory University School of Medicine, Atlanta, GA, USA); Ana Fernandez-Sesma (Icahn School of Medicine at Mount Sinai, New York, New York, USA); Alessandro Sette (La Jolla Institute for Immunology, La Jolla, CA, USA); Ken Stuart (Seattle Children's Research Institute, Seattle, WA, USA); Alkis Togias (NIAID, NIH, Bethesda, MD, USA); John S Tsang (NIAID and Center for Human Immunology (CHI), NIH, Bethesda, MD, USA)
